# A single-cell trajectory atlas of striatal development

**DOI:** 10.1101/2022.11.18.517098

**Authors:** Ashley G. Anderson, Ashwinikumar Kulkarni, Genevieve Konopka

**Affiliations:** Department of Molecular and Human Genetics, Baylor College of Medicine, Houston, TX; Department of Neuroscience, UT Southwestern Medical Center, Dallas, TX, 75390-9111, USA

## Abstract

The striatum integrates dense neuromodulatory inputs from many brain regions to coordinate complex behaviors. This integration relies on the coordinated responses from distinct striatal cell types. While previous studies have characterized the cellular and molecular composition of the striatum using single-cell RNA-sequencing at distinct developmental timepoints, the molecular changes spanning embryonic through postnatal development at the single-cell level have not been examined. Here, we combine published mouse striatal single-cell datasets from both embryonic and postnatal timepoints to analyze the developmental trajectory patterns and transcription factor regulatory networks within striatal cell types. Using this integrated dataset, we found that dopamine receptor-1 expressing spiny projection neurons have an extended period of postnatal development with greater transcriptional complexity compared to dopamine receptor-2 expressing neurons. Moreover, we found the transcription factor, FOXP1, exerts indirect changes to oligodendrocytes. These data can be accessed and further analyzed through an interactive website (https://cells-test.gi.ucsc.edu/?ds=mouse-striatal-dev).

## Introduction

The striatum is a highly conserved forebrain structure important for regulating a wide range of motor and cognitive behaviors (Gerfen and Surmeier 2011). This region receives dense glutamatergic and neuromodulatory inputs from several brain regions, including the cortex, thalamus, and substantia nigra (Gerfen and Surmeier 2011). These diverse inputs are integrated and propagated to downstream basal ganglia nuclei via distinct classes of striatal GABAergic spiny projection neurons (SPNs), interneurons, and glial cell types. Disruption of striatal cell types has been observed across several neurodevelopmental and neurodegenerative disorders, including autism spectrum disorders (ASD) and Huntington’s disease (HD) (Crittenden and Graybiel 2011, Fuccillo 2016). Uncovering the molecular mechanisms regulating striatal cell type development in the brain is therefore an important step towards identifying mechanisms altered in disease states to ultimately improve therapeutics.

High throughput single-cell RNA-sequencing (scRNA-seq) technology has advanced our understanding of the cellular composition and molecular characterization of the brain (Kulkarni, Anderson et al. 2019). In the striatum specifically, recent scRNA-seq studies have led to important insights into striatal cellular composition and further unraveled the molecular differences between the principal striatal spiny projection neurons and aspiny interneuron subtypes (Chen, Friedman et al. 2017, Munoz-Manchado, Bengtsson Gonzales et al. 2018, Saunders, Macosko et al. 2018, Zeisel, Hochgerner et al. 2018, Martin, Calvigioni et al. 2019, Anderson, Kulkarni et al. 2020, Stanley, Gokce et al. 2020). SPNs and interneurons are derived from separate progenitor pools from either the lateral ganglionic eminence (LGE) or medial and caudal ganglionic eminence (MGE, CGE), respectively. SPNs are classically divided into neurons that express dopamine receptor-1 receptor (D1) and project along the *direct* pathway (dSPNs) or SPNs that express dopamine receptor-2 (D2) and project along the *indirect* pathway (iSPNs) (Gerfen and Surmeier 2011). scRNA-seq studies have found more diversity among SPN subtypes than previously appreciated, including a small population of neurons that express either D1 or D2 receptors but have distinct molecular profiles from canonical SPNs (“eccentric” SPNs, or eSPNs) (Saunders, Macosko et al. 2018). This population was masked by previous studies relying on fluorescent-reporter-driven techniques (Valjent, Bertran-Gonzalez et al. 2009) to identify and separate dSPNs versus iSPNs for molecular characterization followed by bulk RNA-sequencing approaches, showing the importance of single-cell methodologies (Heiman, Schaefer et al. 2008, Lobo, Covington et al. 2010, Maze, Chaudhury et al. 2014). Striatal scRNA-seq studies have also shed light on the cell type specific molecular changes that occur upon disrupting genes important for striatal development, such as *FOXP1*, a gene strongly associated with autism and intellectual disability in humans (Anderson, Kulkarni et al. 2020). Though highly expressed in both dSPNs and iSPNs, deletion of *Foxp1* severely affected iSPNs and significantly reduced that cellular population specifically (Anderson, Kulkarni et al. 2020). Striatal interneurons make up ~5% of the striatal neuron population and are largely divided into subtypes that include *Pvalb-expressing, Sst/Npy/Nnos-expressing, Calb2*-expressing, and *Th*-expressing groups (Munoz-Manchado, Bengtsson Gonzales et al. 2018, Tepper, Koos et al. 2018). Singlecell studies of striatal interneurons have found interneurons subgroups that are molecularly discrete (i.e. *Npy^+^/Sst^+^, Chat^+^, Th^+^, Npy^+^/Sst^-^, Cck^+^*) or display continuous gradients of gene expression (*Pvalb*) (Munoz-Manchado, Bengtsson Gonzales et al. 2018). While these studies have furthered our understanding of striatal cellular and molecular development, each study was performed at a single time point and the developmental trajectory of striatal cell types spanning development remains incomplete. A recent single-cell study has made some progress along these lines by profiling cells from the human fetal lateral ganglionic eminence from 7 to 11 post conceptual weeks and identified key transcription factors important for governing D1 and D2 lineage specification, highlighting the benefit of trajectory-level analyses (Bocchi, Conforti et al. 2021).

To identify the key molecular mechanisms spanning striatal development, we combine previously published single-cell or single-nuclei RNA-sequencing datasets at distinct embryonic and postnatal timepoints in the mouse brain to build a striatal cell type-specific developmental trajectory map (Chen, Friedman et al. 2017, Munoz-Manchado, Bengtsson Gonzales et al. 2018, Saunders, Macosko et al. 2018, Zeisel, Hochgerner et al. 2018, Martin, Calvigioni et al. 2019, Anderson, Kulkarni et al. 2020, Stanley, Gokce et al. 2020). From this integrated dataset, we investigate the trajectory pattern and gene regulatory networks within both neuronal and glial populations. We find that dSPNs and iSPNs diverge in their postnatal developmental trajectory, with dSPNs exhibiting a prolonged developmental window compared to iSPNs. Moreover, dSPNs exhibit greater transcriptional complexity compared to iSPNs. We further show how interneuron subtypes and oligodendrocytes change their molecular composition over development. Moreover, we show that FOXP1 may indirectly alter oligodendrocyte maturation via SPN-specific disruption. We created an interactive website to easily access and further analyze these datasets. This resource is an important step towards compiling single-cell data from across labs and methodologies to further our understanding of neural development.

## Results

### Combined striatal single-cell datasets across development

To build a single-cell developmental trajectory map of striatal cell types, we integrated previously published striatal single-cell datasets from seven studies that collected data from mouse brain at different timepoints during striatal development (**Figure 1A**). These data include single cells from medial and lateral ganglionic eminences and mature striatal tissue between the ages of embryonic day (E) E11.5-E17.5 (Chen, Friedman et al. 2017) (C17, 225 cells), postnatal day (P) P9 (Anderson, Kulkarni et al. 2020) (A20, 14,467 cells), P12-P30 (Zeisel, Hochgerner et al. 2018) (Z18, 31,836 cells), P22-P28 (Munoz-Manchado, Bengtsson Gonzales et al. 2018) (Dataset A, MA18: 1,122 cells and Dataset B, MB18: 3,417 cells), P35-47 (Stanley, Gokce et al. 2020) (S20, 1,207 cells), P60-P70 (Saunders, Macosko et al. 2018) (S18, 75,469 cells), and P56-112 (Martin, Calvigioni et al. 2019) (M19, 768 cells). The number of genes detected per dataset was related to the number of cells sequenced, with more genes present in datasets with fewer cells that were more deeply sequenced using Smart-seq2 (C17, S20, and M19, **Figure S1A**).

**Figure 1.**
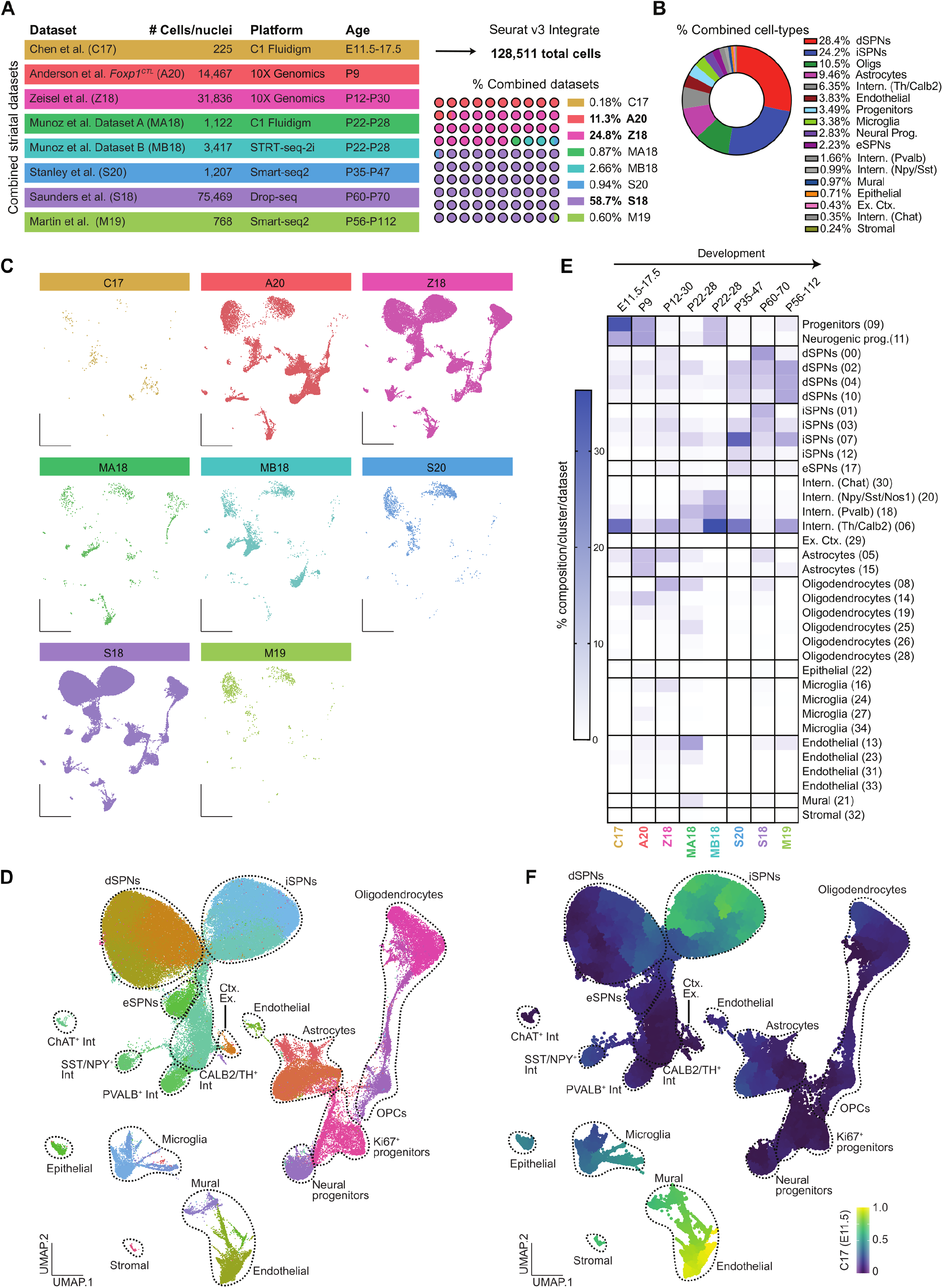
Integrated striatal single-cell datasets across timepoints and single-cell methodologies. **(A)** Table describing the published datasets integrated in our analysis and the percent composition of each study to the combined dataset. **(B)** The percent composition of cell types within the combined dataset. **(C)** UMAP plots showing where cells from each dataset clustered in the combined analysis. **(D)** UMAP of combined dataset colored by cluster affiliation and annotated with cell type identification. **(E)** The percent contribution of cells per cluster (35 total) for each dataset ordered by developmental time. **(F)** Pseudotime analysis using Monocle3 of the combined dataset plotted with UMAP coordinates.

The combined dataset resulted in 128,511 total cells with 35 unique clusters (**Figure S1C**). Three datasets across postnatal development contributed greater than 95% of cells to the combined analysis (**Figure 1A**). dSPNs (28.4%) and iSPNs (24.2%) comprised ~52.6% of the total dataset, followed by oligodendrocytes (10.5%), astrocytes (9.5%), and interneurons (6.35%) (**Figure 1B**). No clusters were unique to a given dataset (**Figure 1C and Figure S1C**), with cells clustering primarily by cell type identity (**Figure 1D and Figure S1D**). The number of genes were greater in neuronal cell types (**Figure S1B**) or across neuronal clusters (**Figure S1C**) as seen in previous single-cell studies of brain tissue (Hodge, Bakken et al. 2019). We found that the percentage of cells from the embryonic and early postnatal timepoints were more abundant in the progenitor and neural progenitor clusters compared to P12-P112 timepoints (**Figure 1E).** Four datasets used cellular isolation methods to enrich for distinct cell types, interneurons for MA18 and MB18 (Munoz-Manchado, Bengtsson Gonzales et al. 2018) and SPNs for M19 and S20 (Martin, Calvigioni et al. 2019, Stanley, Gokce et al. 2020), which is observed in the percent composition of cell types within these studies (**Figure 1E**).

To examine the differentiation trajectory pattern of the combined dataset, we used Monocle3 (Cao, Spielmann et al. 2019) to organize cells along a pseudotime scale by setting the root as the biologically earliest cells (E11.5). We then projected their pseudotime values onto UMAP coordinates (**Figure 1F**). Using this method, we observed the most dynamic changes in cellular trajectory patterns within three cell types: SPNs, microglia, and vascular cells (**Figure 1F**). Within the SPN population, a distinct change in trajectory pattern was observed between dSPNs and iSPNs, with iSPNs progressing faster along the differentiation trajectory compared to dSPNs. These findings suggest dSPNs and iSPNs have distinct developmental trajectory patterns.

### Extended period of gene expression dynamics in dSPNs relative to iSPNs

To further study the pseudotime trajectories of dSPNs versus iSPNs, we isolated SPNs from the combined dataset, using clusters 0, 2, 4, 10 for dSPNs and 1, 3, 7, 12 for iSPNs (**Figure 1E**). We used PHATE (Moon, van Dijk et al. 2019) to perform a pseudotime analysis only on SPNs (**Figure 2A**). Similar to the results found using Monocle3 (Cao, Spielmann et al. 2019) on all cells, iSPNs were farther along in pseudotime compared to dSPNs (**Figure 2A-B**). To quantitatively compare the trajectory dynamics between dSPNs and iSPNs we used cellAlign (Alpert, Moore et al. 2018) to compare single-cell pseudotime trajectories. The outputs of this analysis are a global alignment-based dissimilarity matrix and a pseudotime shift score indicating differences between pseudotime values (**Figure 2C**). Using this method, we observed a distinct pseudotime shift between iSPNs and dSPNs, indicating that faster gene expression dynamics occur within iSPNs relative to dSPNs (**Figure 2C**). This temporal lag in dSPN development hints at an extended period of gene expression dynamics during dSPN maturation (**Figure 2C**). This change in pseudotime dynamics is observed across each dataset when plotting the expression of dSPN markers (*Drd1, Tac1*, **Figure 2D**) or iSPN markers (*Penk, Drd2*, **Figure 2E**) across pseudotime. We note that the embryonic C17 dataset has low signal for dSPNs. Therefore, we used the early postnatal (P9) cells to set the pseudotime trajectory root and observed the same pseudotime shifts, suggesting that this difference in relative gene expression dynamics occurs during postnatal development (**Figure S2**).

**Figure 2.**
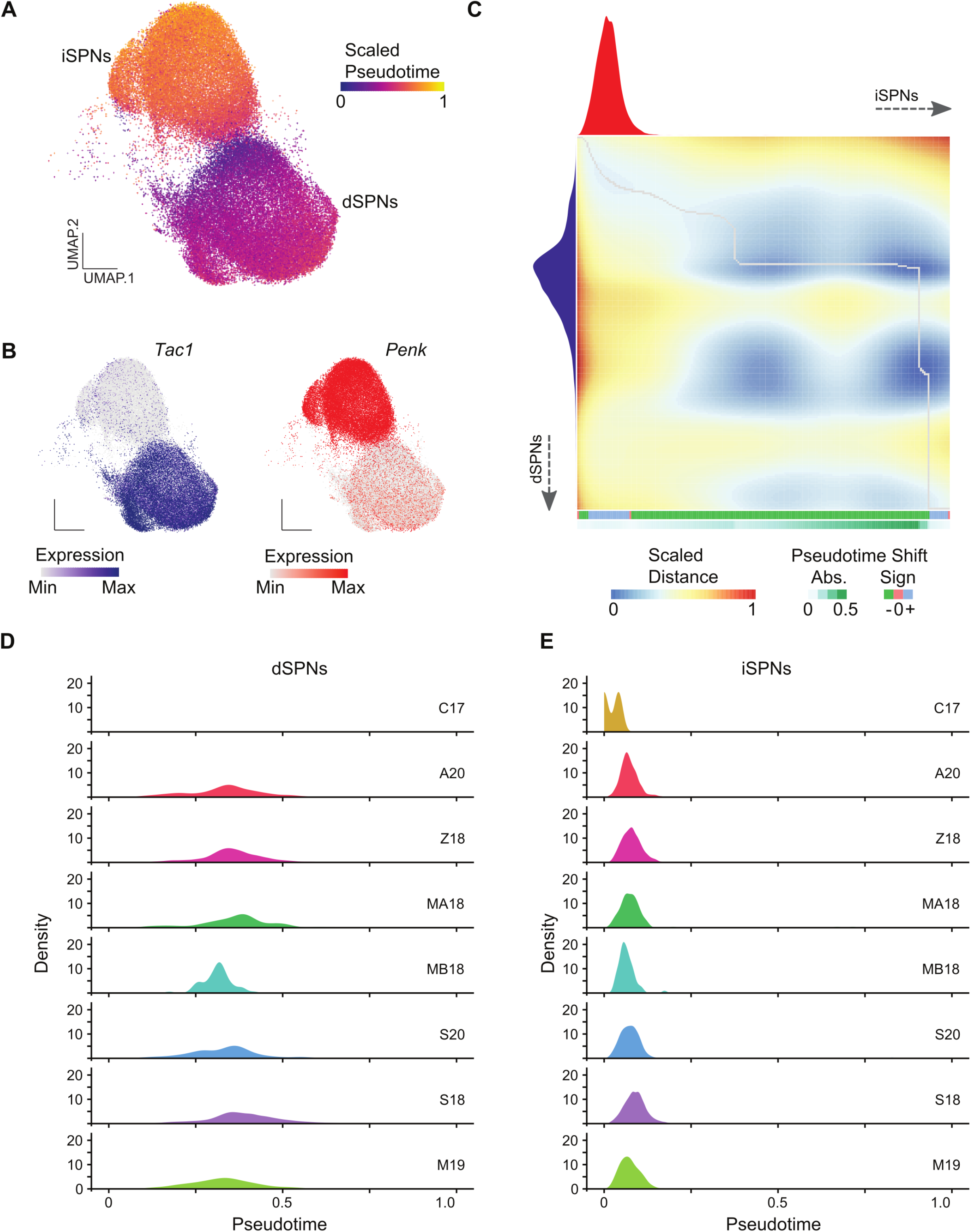
Comparing pseudotime trajectory dynamics between dSPNs and iSPNs. **(A)** UMAP plot colored by PHATE pseudotime scale with **(B)** feature plots showing the expression of dSPN marker (*Tac1*) and iSPN marker (*Penk*). **(C)** Dissimilarity matrix and global alignment of pseudotime trajectories between dSPNs (x-axis) and iSPNs (y-axis) with pseudotime shifts labelled below. **(D)** Plots of *Tac1* (dSPNs) or **(E)** *Penk* (iSPN) expression across pseudotime separated by dataset.

### dSPNs have more discrete transcriptional networks

We next used a gene regulatory network (GRN) analysis to identify key transcription factors (TFs) involved in dSPN and iSPN development (**Figure 3**). dSPNs and iSPNs have several shared hub TFs including *Foxp1, Myt1l, Meis2*, and *Csde1*. We also identified hub TFs unique to each subpopulation. dSPNs unique hub TFs included *Sox11, Bcl11b, Ybx1*, and *Ebf1* (**Figure 3A**). iSPNs unique hub TFs included *Rarb, Nr1d1*, and *Tef* (**Figure 3B**). We observed that dSPNs had more discreet transcriptional networks, compared to iSPNs whose hub genes were more interconnected. Moreover, the dSPNs TF hub genes were enriched with markers of early-born neurons, including *Sox4* and *Sox11* (**Figure 3A**). These results suggest that dSPNs have more transcriptional complexity compared to mature iSPNs.

**Figure 3.**
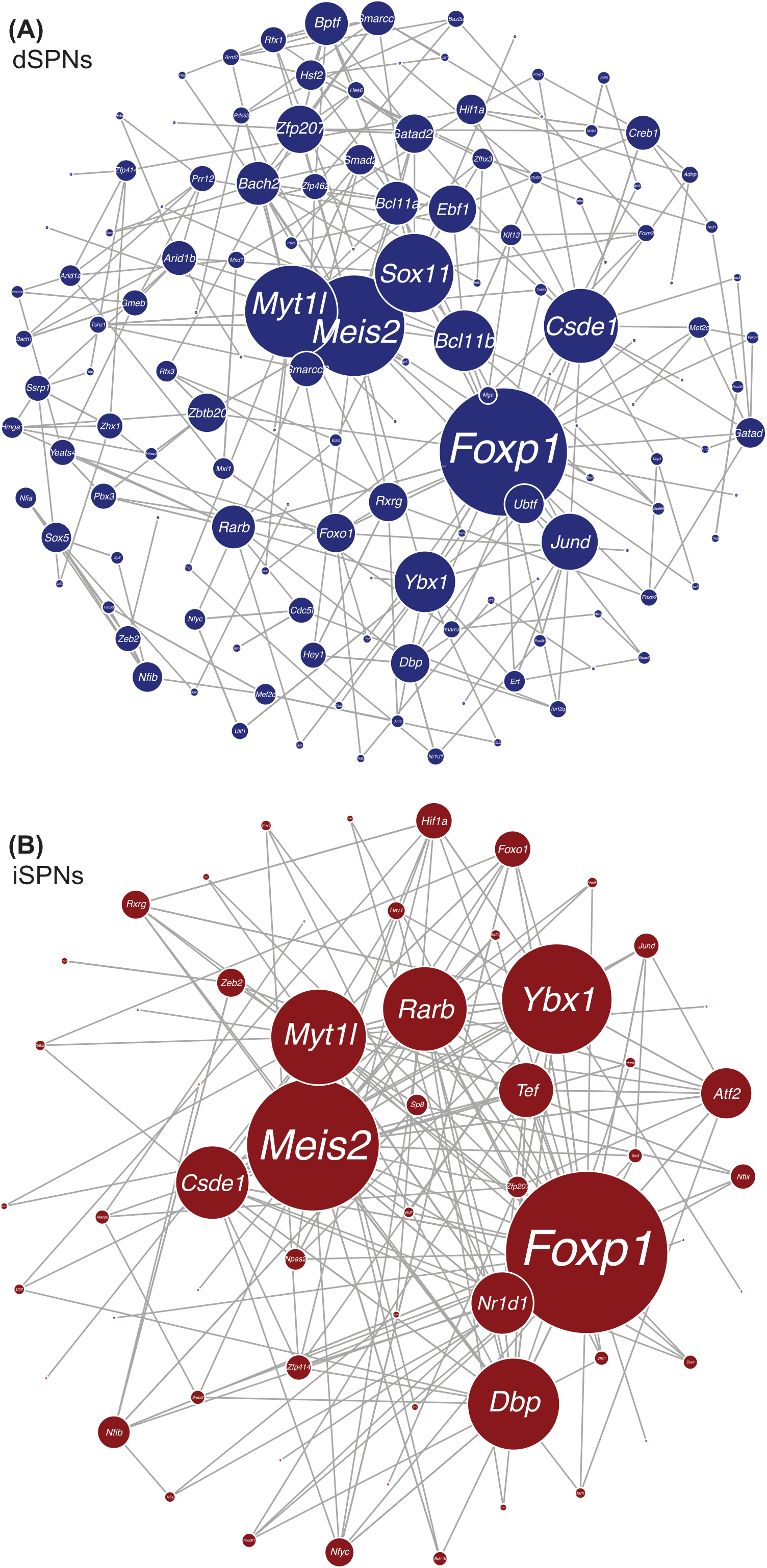
Gene regulatory analysis of dSPN and iSPNs. Visualization of network transcription factors in dSPNs **(A)** or iSPNs **(B)**. Gray lines between hubs indicate degree of interconnectivity.

### Interneuron pseudotime trajectories over striatal development

Several interneuron subtypes populate the striatum and are critical for striatal function. We isolated both interneuron and neural progenitor clusters from the integrated dataset to analyze the pseudotime trajectory pattern of these interneuron subtypes (**Figure 4A**). We identified interneurons by key molecular markers including *Chat* (cholinergic interneurons), *Npy* (Neuropeptide Y), *Nos1* (Nitric oxide synthase 1), *Sst* (somatostatin), *Pvalb* (Parvalbumin), *Th* (Tyrosine hydroxylase), and *Calb1* (Calbindin 1) (**Figure 4B**). Several of these markers colocalize in the same cells (*Nos1, Npy, Sst*), whereas *Th, Pvalb, Calb1*, and *Chat* interneuron clusters were distinct. Using PHATE (Moon, van Dijk et al. 2019), we found distinct differences in the pseudotime differentiation trajectory of interneuron subtypes (**Figure 4C**). *Chat* and *Nyp/Sst* interneurons were further along in pseudotime compared to the other subtypes, followed by *Pvalb, Th*, and *Calb2* expressing interneurons. (**Figure 4C**). We next plotted the expression of key markers of the progenitor state (*Sox4, Sox11, Dlx2, Mki67, Ascl1*), interneuron markers, and the TFs highly associated with interneuron development (*Sox2/5/6/9, Pax6, Lhx2, Nkx2.1, Etv1, Lhx6/8*, and *Nr2f2*) (**Figure 4D**) (Lim, Mi et al. 2018). As expected, peak expression of progenitor markers occurred early in pseudotime, whereas interneuron markers peaked later in pseudotime with little to no overlap. We also saw that *Lhx6* and *Lhx8* expression peaked along the scaled pseudotime after *Nkx2.1* expression, since both are downstream of *Nkx2.1*. Moreover, *Lhx6* is critical for *Pvalb* and *Sst/Npy/Nos1* interneuron specification. *Lhx8* increased over pseudotime following the same trend as *Chat*, an expected finding given that *Lhx8* is important for *Chat* interneuron development and function. Interestingly, we found a bimodal pseudotime pattern of many TFs associated with interneuron development, suggesting successive or distinct waves of interneuron development. This patterning could potentially represent regional differences from interneurons derived from different subregions within the medial or caudal GE, since cell types from both regions have unique cellular trajectories (Lee, Rhodes et al. 2022). These results indicate that interneuron subtypes develop along distinct trajectory patterns and provide a rich resource for researchers to further investigate molecular development of striatal interneurons.

**Figure 4.**
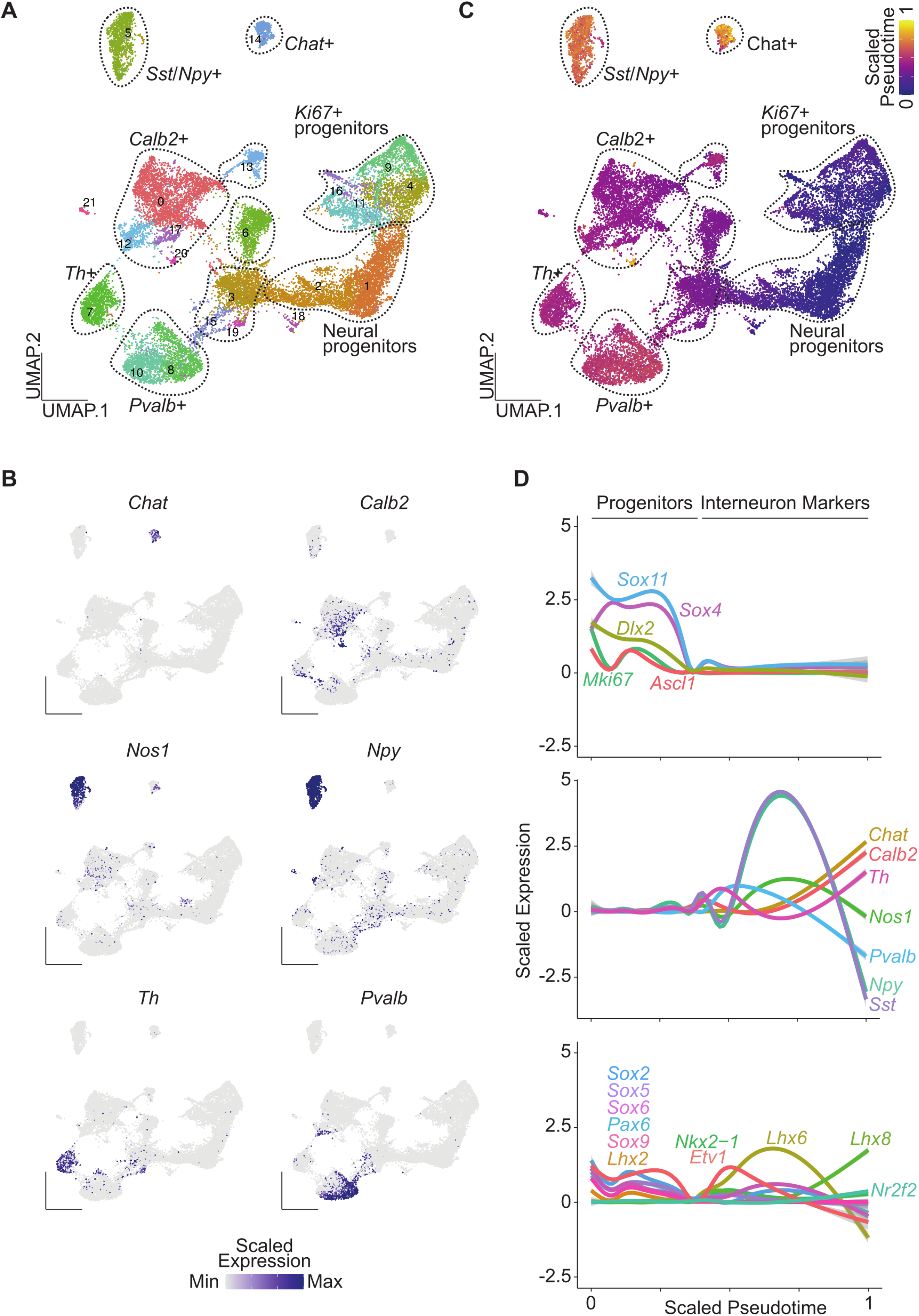
Pseudotime analysis of striatal interneuron subtypes. **(A)** Annotated UMAP clusters of interneuron and neural progenitor clusters isolated from the integrated dataset. **(B)** Scaled expression of genes enriched in distinct interneuron populations. **(C)** UMAP of cells colored by PHATE pseudotime values. **(D)** Expression of progenitor markers (top panel), interneuron subtype markers (middle panel), and key transcription factor important for interneuron development (bottom panel) across pseudotime.

### Oligodendrocyte pseudotime trajectories over striatal development

We next wanted to examine the developmental trajectory of the second most abundant cell type within this integrated dataset, oligodendrocytes. We isolated both the oligodendrocyte and progenitor clusters from the integrated dataset and identified clusters for oligodendrocyte precursors (OPC, cluster 1), committed oligodendrocyte precursors (COP, cluster 11), newly-formed oligodendrocytes (NFOL, cluster 12), myelin-forming oligodendrocytes (MFOL, clusters 0, 2, 8 and 13), mature oligodendrocytes (MOL, cluster 4) along with progenitors (PROG, clusters 3, 7, 9, 10 and 15) (**Figure 5A**). Using PHATE to obtain pseudotime trajectory values for each cell, we found distinct trajectory originating from progenitors (*Mki67*^+^) to clusters enriched for markers of mature oligodendrocytes (*Klk6^+^, Apod^+^*) progressing through OPCs, COPs, NFOL and MFOL (**Figure 5B**). OPCs were marked by the expression of gene markers such as *Pdgfra* and *Cspg4* (**Figure 5C**). Genes previously associated with astrocytes or radial glia (*Tmem100*) also appeared to be enriched in OPCs consistent with the origin of OPCs from radial glia-like cells and their ability to generate astrocytes in an event of injury (Dimou and Gotz 2014, Marques, Zeisel et al. 2016) (**Figure 5C**). COPs were distinct from OPCs and expressed *Neu4* and genes involved in keeping oligodendrocytes undifferentiated such as *Bmp4* (Samanta and Kessler 2004, Marques, Zeisel et al. 2016) (**Figure 5C**). NFOLs expressed genes involved in oligodendrocyte differentiation such as *Tcf7l2* (Ye, Chen et al. 2009, Marques, Zeisel et al. 2016) (**Figure 5C**). Both COPs and NFOLs also showed expression of genes involved in migration such as *Tns3* (Marques, Zeisel et al. 2016) (**Figure 5C**). MFOLs expressed genes such as *Mal* and *Opalin* known to be responsible for myelin formation whereas MOLs expressed late oligodendrocyte differentiation genes (*Klk6, Apod*) including genes enriched in myelinating cells (*Pmp22*) (Cahoy, Emery et al. 2008, Marques, Zeisel et al. 2016) (**Figure 5C**). These findings show that striatal oligodendrocytes have distinct subtypes with unique gene expression and trajectory profiles.

**Figure 5.**
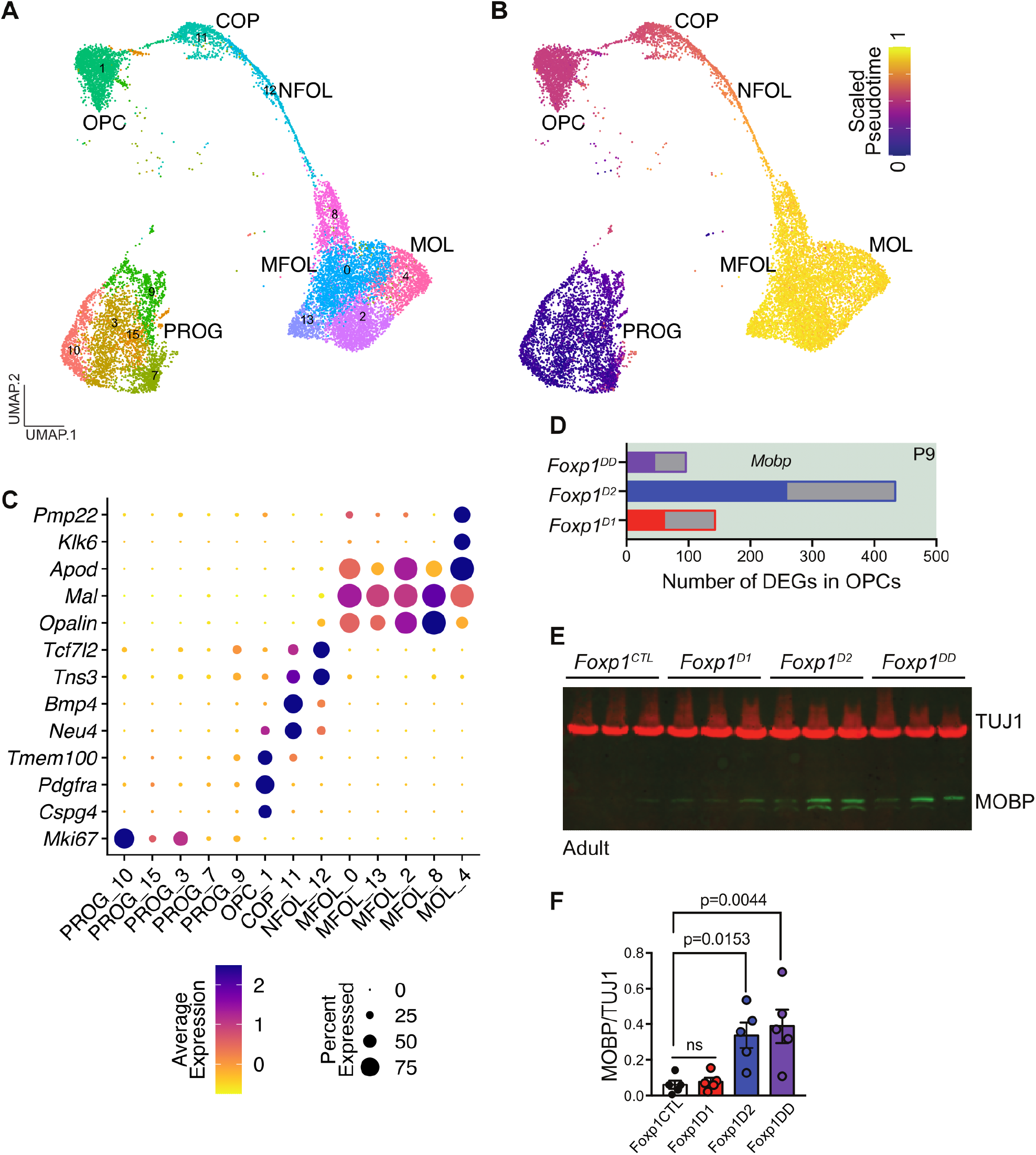
Pseudotime analysis of oligodendrocytes across striatal development and with deletion of *Foxp1*. **(A)** Annotated UMAP of oligodendrocyte progenitor cells (OPCs), committed oligodendrocyte precursors (COPs), newly-formed oligodendrocytes (NFOL), myelin-forming oligodendrocytes (MFOL), mature oligodendrocytes (MOL) and progenitor clusters from the combined dataset. **(B)** Cells colored by PHATE pseudotime values. **(C)** Scaled expression of genes specific to each oligodendrocyte subtype. **(D)** Number of differentially expressed genes (DEGs) in OPCs from P9 dataset with deletion of *Foxp1* from dSPNs, iSPNs, or both. Filled bars are upregulated DEGs (such as *Mobp*) and grey bars are downregulated DEGs. **(E)** Western blot of TUJ1 (housekeeping gene) and MOBP (oligodendrocyte marker) in striatal tissue in *Foxp1^CTL^, Foxp1^D1^, Foxp1^D2^*, and *Foxp1^DD^* mice and **(F)** quantification of MOBP levels relative to TUJ1 across genotypes (N=5/genotype). Data is represented as mean ± SEM. P-values determined using a one-way ANOVA with Dunnett’s multiple comparison.

### Deletion of *Foxp1* upregulates oligodendrocyte marker MOBP in striatum

To better understand the non-cell autonomous effects of disrupting a key transcription factor in striatal SPN development (**Figure 3**), we performed a differential gene expression analysis within oligodendrocytes in P9 striatal cells with *Foxp1* deleted from either dSPNs (*Foxp1^D1^*), iSPNs (*Foxp1^D2^*), or both (*Foxp1^DD^*). We found both upregulated and downregulated DEGs in oligodendrocytes across all genotypes, but more DEGs were observed when *Foxp1* was deleted specifically from iSPNs (**Figure 5D**). The mature oligodendrocyte marker, *Mobp*, was upregulated in *Foxp1^D2^* samples and we confirmed this finding at the protein level (**Figure 5F**). While deletion of *Foxp1* in iSPNs was shown to have non-cell-autonomous effects on dSPNs (Anderson, Kulkarni et al. 2020), we now show that loss of *Foxp1* in iSPNs also exerts non-cell-autonomous effects on oligodendrocytes in the striatum.

## Discussion

The striatum is a hub for propagating signals from multiple brain regions to modulate complex learning and motor behaviors. Here, we have developed a single-cell transcriptome resource with the goal of increasing understanding of striatal molecular development at cellular resolution. We have developed an interactive website that integrates previously published striatal single-cell datasets across timepoints and technological modalities. This resource can also be expanded to include additional datasets and can be easily navigated by bench scientists.

Using this integrated striatal single-cell dataset, we analyzed trajectory information for the main neuronal cell types (dSPNs, iSPNs, and interneurons) and one major glial cell type (oligodendrocytes) of the striatum. Our findings also suggest that dSPNs have a period of prolonged development and discrete transcriptional states compared to iSPNs. In the study of human embryonic striatal scRNA-seq, a previous study found that dSPNs had slower differentiation kinetics compared to iSPNs and dSPNs had a greater number of transcriptionally distinct clusters (Bocchi, Conforti et al. 2021). These data are in line with our findings that dSPNs have greater transcriptional complexity compared to iSPNs. This is interesting given the enrichment of dSPNs in distinct neurochemical compartments of the striatum, known as the striosome (or “patch”), compared to iSPNs. Striosomes receive dense dopaminergic innervation from the VTA and substantia nigra. This innervation becomes more dense over postnatal development, which might play a role in the different trajectory patterns and transcriptional states observed between dSPNs and iSPNs in our analysis.

The striatum contains a substantial population of oligodendrocytes and these cells likely constitute the increased amount of myelination that occurs postnatally on axon tracts targeting and passing through the striatum. Oligodendrocytes are responsible for generating myelin sheaths for the optimization of signal conductance, maturation, survival, and regenerative properties of axons. They are also vulnerable to dysfunction in numerous disorders, including ASD and HD. For example, oligodendrocyte density is increased within HD post-mortem striatum compared to healthy controls. A mouse model of Timothy syndrome, a severe congenital syndrome associated with autism and caused by mutations in an L-type voltage-gated Ca+ channel (Cav1.2), exhibits accelerated oligodendrocyte development and myelination in the striatum (Cheli, Santiago Gonzalez et al. 2018). How oligodendrocytes mature in the striatum over development at the single-cell level is unknown. We found that striatal oligodendrocytes have a distinct lineage with different developmental stages.

Non-neurons, including oligodendrocytes, can send and receive signals to neurons. Such interactions are ultimately important for normal development and function of neurons. Single cell genomics can be harnessed to uncover non-cell autonomous effects on gene expression with the alteration of individual genes in specific cell types. Thus, we asked whether manipulation of striatal SPNs might alter non-neuronal populations in the striatum. We examined how deletion of the transcription factor *Foxp1*, a hub transcription factor in our GRN analysis of SPNs, alters the trajectory pattern of striatal oligodendrocytes. We identified non-cell autonomous gene expression changes in oligodendrocytes with deletion of *Foxp1* in dSPNs, iSPNs, or both cell types. Similar to the Timothy syndrome mouse model, we found that loss of FOXP1 specifically in iSPNs enhanced the maturation of oligodendrocytes and significantly increased the mature oligodendrocyte marker MOBP in adult striatum. These findings are just one example of how this resource can be queried to understand the role of individual genes on cell type specific patterns of expression over striatal development in both a cell autonomous and non-cell autonomous manner. Ultimately, this resource should further our understanding of striatal neurobiology at the single-cell level and aid in addressing therapeutic challenges facing neurodevelopmental and degenerative disorders that alter striatal function.

## Materials and Methods

### Integration analysis

First, raw counts, matching cell type and meta information for each of the datasets was downloaded from respective sources. After checking the integrity of the datasets, raw counts for only common protein-coding genes across all the datasets were retained. Data processing and analysis was performed using R. Individual datasets were first filtered following cutoffs mentioned in each published paper (see table below). For dataset(s) with ‘NA’ cutoffs, either the datasets were already filtered and/or mitochondrial genes were already filtered out. Also, genes with no expression in any of the cells and genes from chromosomes X, Y and M were removed. Following the filtering, each dataset was processed through the standard Seurat (v3) pipeline (*NormalizeData, FindVariableFeatures, ScaleData, FindNeighbors, RunUMAP, FindClusters*) regressing for the number of UMIs and percent mitochondrial content (https://satijalab.org/seurat/archive/v3.0/pbmc3k_tutorial.html) (Stuart, Butler et al. 2019). Seurat objects for each of the datasets were then combined using Seurat’s integration (https://satijalab.org/seurat/archive/v3.0/integration.html) (Stuart, Butler et al. 2019) approach (*FindIntegrationAnchors, IntegrateData*) with 30 principal components. Data were clustered (*FindNeighbors, FindClusters*) using the original Louvain algorithm with a resolution of 0.8 and the clusters were visualized with Uniform Manifold Approximation and Projection (UMAP) (Becht, McInnes et al. 2018, Kulkarni, Anderson et al. 2019) in two dimensions (*RunUMAP*) for a total of 128,511 single cells or nuclei from the mouse striatum. Gene markers enriched for each cluster were identified using ‘*FindAllMarkers*’. Clusters were then annotated using the ‘*LabelTransfer*’ approach from Seurat (https://satijalab.org/seurat/archive/v3.0/integration.html) (Stuart, Butler et al. 2019) using cell types defined in S18 as reference.

### Pseudotime trajectory analysis for all cell types

The integrated Seurat object with all cell types for all datasets was converted into a Monocle (v3) compatible object using the ‘*as.cell_data_set*’ command. The Monocle object was then pre-processed (*cluster_cells, learn_graph*) using the standard Monocle pipeline (Trapnell, Cacchiarelli et al. 2014, Cao, Spielmann et al. 2019) (https://cole-trapnell-lab.github.io/monocle3/docs/trajectories/). Further, E11.5 cells from C17 were selected as the root population for performing pseudotime trajectory analysis (*order_cells*). UMAP plots colored by scaled pseudotime values were then generated.

| Dataset | # UMI cutoff | % Mito cutoff | # Cells |
| --- | --- | --- | --- |
| C17 | 3,000,000 | NA | 225 |
| A20 | ≤ 50,000 | ≤ 10 | 14,467 |
| Z18 | NA | ≤ 10 | 31,836 |
| MA18 | ≤ 20,000 | NA | 1,122 |
| MB18 | ≤ 20,000 | NA | 3,417 |
| S20 | NA | NA | 1,207 |
| S18 | $\leq 40,000$ | $\leq 10$ | 75,469 |
| M19 | NA | NA | 768 |
| **TOTAL** |  |  | 128,511 |

### SPN sub-clustering and pseudotime trajectory analysis

Cell barcodes corresponding to clusters annotated as SPNs (dSPNs, iSPNs and eSPNs) were then used to subset the SPN population from all the cells. Using raw counts corresponding to the identified SPN population for all datasets were then used to run the Seurat integration approach (*FindIntegrationAnchors, IntegrateData*) to identify sub-populations among major SPN categories [https://satijalab.org/seurat/archive/v3.0/integration.html) (Stuart, Butler et al. 2019). Using 30 principal components, SPNs were clustered (*FindNeighbors*, *FindClusters*) using the original Louvain algorithm with a resolution of 0.8 and clusters were visualized using UMAP (Becht, McInnes et al. 2018, Kulkarni, Anderson et al. 2019). Using the clustering information for SPNs, subsets for dSPNs and iSPNs were further created. The dSPN and iSPN subsets were then subjected to pseudotime trajectory analysis using PHATE (Moon, van Dijk et al. 2019). First, loom objects corresponding to dSPN and iSPN subsets were exported. Using loom objects as input and *scaNpy’s* python implementation of PAGA/PHATE (Moon, van Dijk et al. 2019) (https://scanpy-tutorials.readthedocs.io/en/latest/paga-paul15.html, https://scanpy.readthedocs.io/en/stable/generated/scanpy.external.tl.phate.html), pseudotime trajectories were computed using C17 cells as root populations. A UMAP visualization of cells colored by scaled diffusion pseudotime (DPT) (Haghverdi, Buttner et al. 2016) was also generated. Pseudotime information for dSPNs and iSPNs was then used to align the trajectories using the ‘*cellAlign*’ approach (Alpert, Moore et al. 2018) (https://github.com/shenorrLab/cellAlign) and data were visualized using a heatmap accompanied with pseudotime densities. A similar approach was also used to perform sub-clustering and trajectory analysis of SPNs populations using cells from the P9 dataset as root population (**Figure S2**).

### Gene regulatory network analysis for SPNs

A list of mouse transcription factors (TFs) was obtained from a mouse tissue transcription factor atlas (Zhou, Liu et al. 2017). A unique list of 471 TFs falling into fetal brain and adult brain tissue categories were retained for gene regulatory network analysis using an *Arboreto* and *grnboost2* based approach (Colquitt, Merullo et al. 2021). First, raw counts corresponding to expressed (446 / 471) TFs was fetched separately for both dSPNs and iSPNs. A gene regulatory network (GRN) was built with raw expression data for dSPNs and iSPNs separately using python implementation of *Arboreto* and *grnboost2* (https://arboreto.readthedocs.io/en/latest/examples.html). The GRN output was then filtered following a previously published approach (Colquitt, Merullo et al. 2021), (https://github.com/bradleycolquitt/songbird_cells/tree/master/grn) to retain the top one percent of the TF-gene interactions, which were then visualized using the *igraph* R package (https://igraph.org/r/).

### Interneurons and oligodendrocytes sub-clustering and pseudotime trajectory analysis

Cell barcodes corresponding to clusters annotated as interneurons (*Pvalb^+^, Sst^+^/Npy^+^, Chat^+^, Calb2^+^/Th^+^*) were then used to subset from all the cells along with *Mki67^+^* progenitors and *Sox4^+^/Sox11^+^* neurogenic progenitors. Raw counts corresponding to the identified subset population for all datasets were then used to run Seurat integration approach (*FindIntegrationAnchors, IntegrateData*) to identify sub-populations among major interneuron categories (https://satijalab.org/seurat/archive/v3.0/integration.html) (Stuart, Butler et al. 2019). Using 30 principal components, interneuron cells were clustered (*FindNeighbors, FindClusters*) using the original Louvain algorithm with a resolution of 0.8 and clusters were visualized using UMAP (Becht, McInnes et al. 2018, Kulkarni, Anderson et al. 2019). The interneuron sub-clustering data were then subjected to pseudotime trajectory analysis using PHATE (Moon, van Dijk et al. 2019). First, loom objects corresponding to interneuron clusters were exported. Using loom objects as input and scaNpy’s python implementation of PAGA/PHATE (Moon, van Dijk et al. 2019), (https://scanpy-tutorials.readthedocs.io/en/latest/paga-paul15.html, https://scanpy.readthedocs.io/en/stable/generated/scanpy.external.tl.phate.html), pseudotime trajectories were computed using cells expressing *Mki67* as root populations. UMAP visualization of cells colored by scaled diffusion pseudotime (DPT) (Haghverdi, Buttner et al. 2016) was also generated. Gene expression patterns for specific sets of progenitor markers, interneuron markers and transcription factors were generated across scaled pseudotime. Similar to interneurons, sub-clustering and pseudotime trajectory analysis was also performed for the oligodendrocyte population including *Mki67^+^* progenitors and a UMAP colored by scaled pseudotime was also generated.

### Mice

All experiments were approved by UT Southwestern IACUC # 2016-101-825. *Foxp1*^flox/flox^ mice were provided by Dr. Haley Tucker and backcrossed to C57BL/6J for at least 10 generations to obtain congenic animals as previously described (Anderson, Kulkarni et al. 2020). *Drd1a-Cre* (262Gsat, 030989-UCD) and *Drd2-Cre* (ER44Gsat, 032108-UCD) mice were obtained from MMRC.

### Protein isolation and immunoblotting

Striatal tissue from adult mice (P56) was harvested as previously described (Anderson, Kulkarni et al. 2020). Briefly, tissue was flash frozen, and protein was extracted using 1X RIPA buffer (750mM NaCl, 250mM Tris-HCl pH7.4, 0.5% SDS, 5% Igepal, 2.5% Sodium deoxycholate, 5mM EDTA, 5mM NaVO4) with fresh protease inhibitor cocktail (10ul/ml), 100mM PMSF (10ul/ml), and 200mM sodium orthovanadate (25ul/ml). Tissue was homogenized using a QIAGEN TissueLyser LT, rotated for 1hr at 4C, and spun down at max speed for 15min. Protein was quantified using a standard Bradford assay and 20ug of protein was loaded into a 10% SDS-Page gel. Protein samples were transferred to a PVDF membrane and then membrane was blocked in a 5% milk TBST solution. The following antibodies were used for immunoblots (IB) experiments: rabbit anti-MOBP (1:2000; Sigma HPA035152) or mouse anti-TUJ1 (1:10,000; Covance MMS-435P). Using an Odyssey infrared imaging system, rectangles were drawn around individual samples in either 800 or 700 IR channels to quantify the intensity signal after setting a background reference rectangle. MOBP signal was normalized to TUJ1 within each sample.

## Acknowledgments

This work was supported by the NIH (NS122920) to A.G.A.; and the Simons Foundation for Autism Research (Award #573689), the James S. McDonnell Foundation 21^st^ Century Science Initiative in Understanding Human Cognition (Scholar Award 220020467) and NIH (MH126481, HG011641, NS115821, MH102603) to G.K.

## Data Availability

These data can be accessed and further analyzed through an interactive website (https://cells-test.gi.ucsc.edu/?ds=mouse-striatal-dev).

---

**Supplemental Figure 1.**
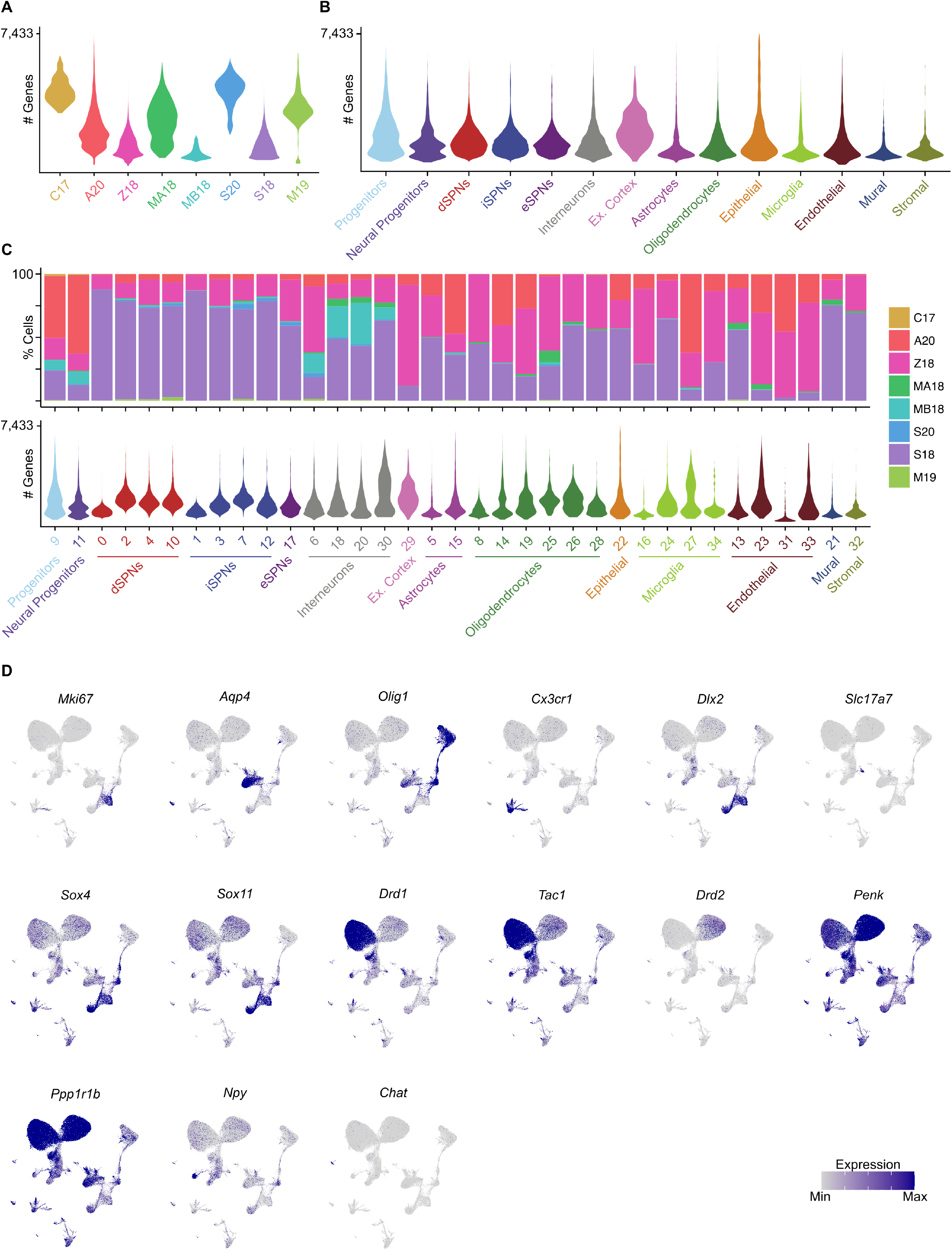
Comparison of genes across datasets and cell types. Comparison of detected genes across **(A)** individual datasets and **(B)** striatal cell types visualized by violin plots. **(C)** Stacked bar plot showing the contribution of cells from individual datasets across the 35 clusters and accompanied by violin plots showing the number of detected genes across striatal cell type clusters. **(D)** Scaled expression of marker genes for each cell type.

**Supplemental Figure 2.**
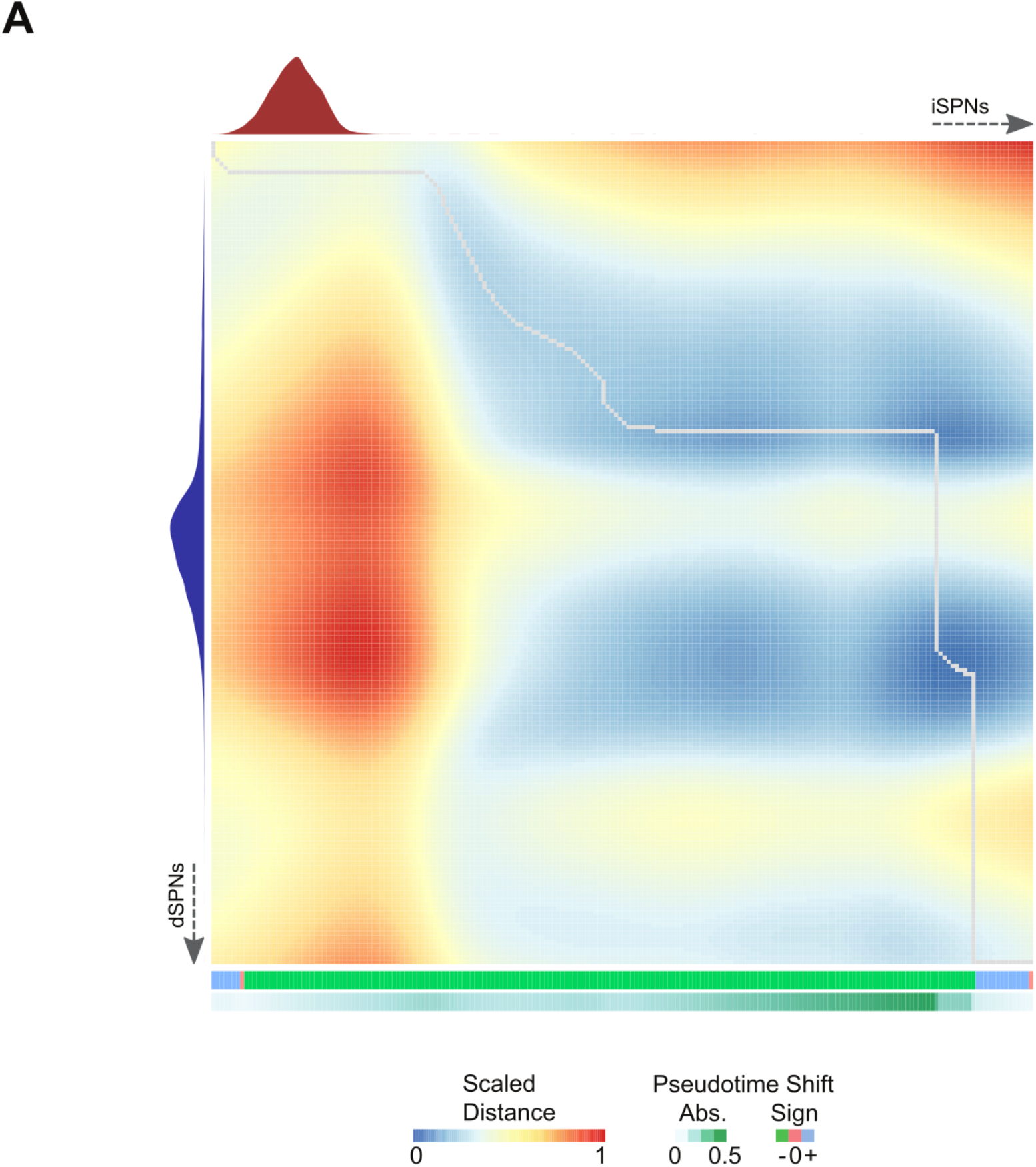
cellAlign analysis of dSPNs and iSPNs. Dissimilarity matrix and global alignment of pseudotime trajectories between iSPNs and dSPNs using cells from the P9 dataset as root.

